# Abstract neural choice signals during action-linked decisions

**DOI:** 10.1101/2020.10.02.323832

**Authors:** Florian Sandhaeger, Nina Omejc, Anna-Antonia Pape, Markus Siegel

## Abstract

Humans can make abstract choices independent of motor actions. However, in laboratory tasks, choices are typically reported with an associated action. Consequentially, knowledge about the neural representation of abstract choices is sparse, and choices are often thought to evolve as motor intentions. Here, we show that in the human brain, perceptual choices are represented in an abstract, motor-independent manner, even when they are directly linked to an action. We measured MEG signals while participants made choices with known or unknown motor response mapping. Using multivariate decoding, we quantified stimulus, perceptual choice and motor response information with distinct cortical distributions. Choice representations were invariant to whether the response mapping was known during stimulus presentation, and they occupied distinct representational spaces from both stimulus and motor signals. Furthermore, their strength predicted decision confidence and accuracy, as expected from an internal decision variable. Our results uncover abstract neural choice signals that generalize to action-linked decisions, suggesting a general role of an abstract choice stage in human decision-making.

## Introduction

Sensory decisions are often linked to an appropriate motor action. This has led to a framework of choices emerging as action intentions (Cisek and Kalaska, 2010), supported by numerous studies showing action-specific choice signals in motor- and premotor areas of the brain (Gold and Shadlen, 2000, 2007; Shadlen and Newsome, 2001; Wang et al., 2019). Compelling evidence favors such an intentional framework over the historic idea of decision-making as a sequential process involving several, successive modules. However, a key component of intelligent behavior is the ability to also make abstract choices when a suitable action is not known in advance (Coallier and Kalaska, 2014; Gold and Shadlen, 2003; Nieder et al., 2020). Any comprehensive account of human decision-making thus has to account for the possibility of abstract choices.

Since most studies use a fixed mapping of perceptual choices (in the following referred to as “choices”) to motor responses, the role of abstraction in sensorimotor decision-making remains elusive. A few notable exceptions, using behavioral tasks with a variable mapping of choices to motor responses, have identified neural representations of abstract choices (Bennur and Gold, 2011; Hebart et al., 2012; Horwitz et al., 2004; Ludwig et al., 2018; Merten and Nieder, 2012, 2013; Minxha et al., 2020; Nieder et al., 2020; Zhou and Freedman, 2019). However, empirical results comparing choice signals in action-linked and action-independent situations are sparse. While some recent work found perceptual choice representations to depend on the ability to plan motor actions (Ludwig et al., 2018; Wang et al., 2019) or response modality (Herding et al., 2017; Wu et al., 2019), other previous evidence suggests at least partially overlapping representations of perceptual choices with specified or unspecified motor actions (Bennur and Gold, 2011).

It is therefore unclear whether, and under which conditions, the same neural representations underlying abstract choice in an action-independent context are also present during choices that are linked to actions. Furthermore, the spatio-temporal dynamics of abstract choice signals are unknown, and it remains unclear whether abstract choice signals constitute an internal decision variable that tracks accumulated evidence. Consequentially, the demonstration of a context-independent, abstract decision variable would be important to confirm predictions of abstraction as an essential stage in perceptual decision-making.

To address this, we investigated human brain activity underlying flexible sensorimotor choices using magnetoencephalography (MEG). The task design and a multivariate analysis framework allowed us to pinpoint abstract neural choice signals in an action-linked as well as in an action-independent context. MEG activity was predictive of participants’ perceptual choices independently of both sensory input and motor behavior. Crucially, a novel metric for the assessment of cross-decoding results enabled us to conclude that abstract choice representations were not only present in both contexts, but indistinguishable between them. Furthermore, choice signals dynamically evolved along the sensorimotor hierarchy and predicted both decision confidence and accuracy, thus exhibiting a hallmark property of an internal decision variable. Our results cast doubt on a purely action-based framework, and suggest a general role for abstraction in sensorimotor decision-making.

## Results

### Behavior in a flexible sensorimotor decision-making task

We recorded MEG in 33 human participants, while they performed variants of a sensorimotor decision-making task (Fig. 1A, see Methods for subsets of participants used for some analyses). In each trial, we presented one of two dynamic random dot stimuli which either contained coherent downwards motion or not (referred to as “signal” and “noise” trials, respectively), and participants judged the presence of coherent motion. To separate stimulus-related neural signals from choice-related signals, we adapted the coherence level in the signal stimulus for each participant such that they performed near threshold. The presence of both correct and error trials then allowed us to identify neural signals associated with the perceptual choice, independent of the physical stimulus, i.e. neural signals that separated correct signal and incorrect noise trials from incorrect signal and correct noise trials. To disentangle choice- and motor response-related signals, we introduced a flexible mapping between perceptual choices and left- or righthand button presses that was cued on a trial-by-trial basis. For half of the trials, the choice-response mapping was revealed before stimulus onset (‘pre-condition’), such that emerging choices could immediately be linked to the appropriate motor response. For the other half (‘post-condition’), we revealed the mapping after stimulus offset, such that participants had to make abstract choices initially, before later selecting their motor response. Participants reported their choices with one of two buttons per choice (inner and outer buttons), thereby additionally indicating their confidence. Participants performed equally well on “pre” and “post” trials (74 % and 73 % correct), neither their sensitivity (d’ = 1.35 and 1.28; *t_25_* = 1.27, *P =* 0.21, two-tailed t-test) nor criterion (C’ = −0.01 and 0.05; *t_25_* = −1.49, *P* = 0.15) were different between tasks, and neither choice was preferentially associated with a particular motor response (50 % “right” responses for both “yes” and “no” choices, *t_25_* = 0.58, *P* = 0.57 and *t_25_* = −0.09, *P* = 0.93, two-tailed t-test). In both task conditions, responses had to be withheld until the fixation point disappeared, and while reaction times (0.74+/-0.23 s, mean +/-standard deviation over participants) were higher in the post-than the pre-condition (Fig, 1B, 0.75 s vs 0.72 s, *F*(1,415) = 7.77, *P* = 0.0056), in noise than in signal trials (*F* = 9.7, *P* = 0.002), and in incorrect than in correct trials (*F* = 34.41, *P* < 10^−8^), they were not significantly different between choices (*F* = 0.81, *P* = 0.37) or responses (*F* = 0.17, *P* = 0.68) (six-way ANOVA including the factors of participant, task condition, stimulus, choice, response and accuracy).

**Fig. 1.**
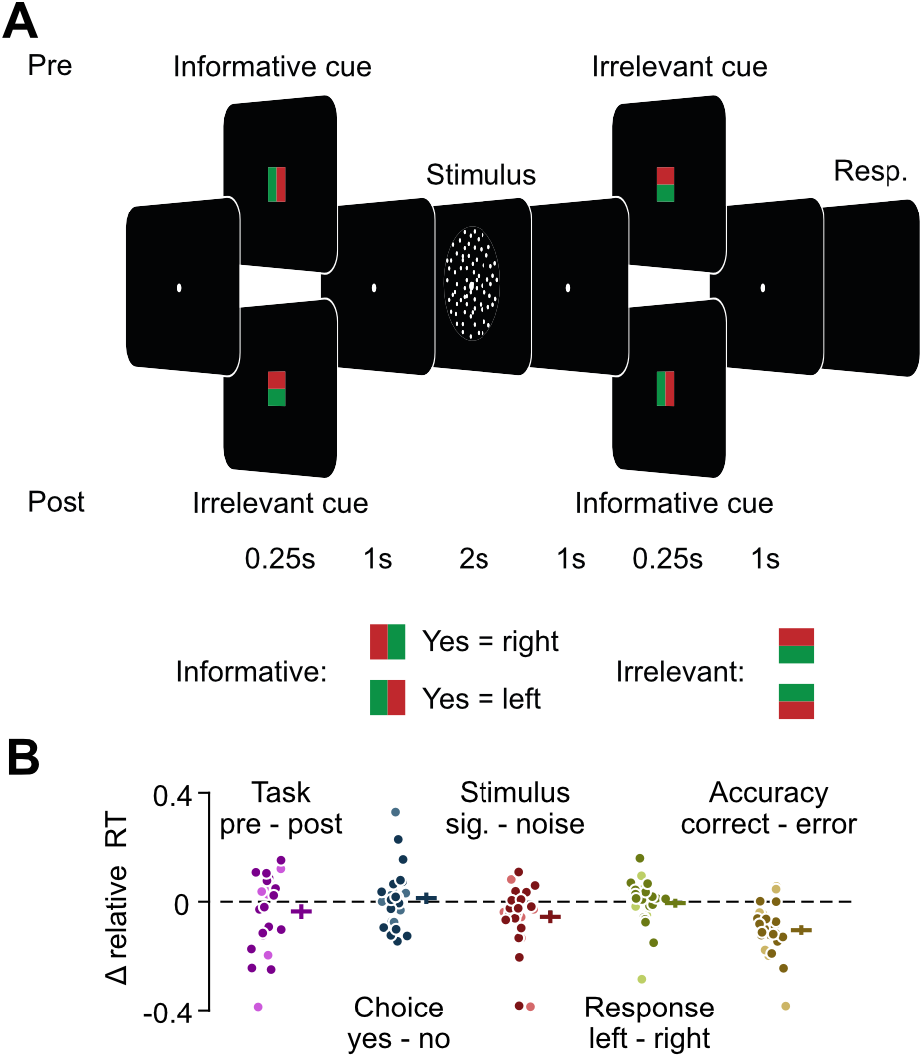
Flexible sensorimotor decision-making task and reaction times. **(A)** In each trial, participants viewed one of two random dot stimuli either containing coherent downwards motion (“signal” trials) or containing only random motion (“noise” trials), and reported the presence of coherent motion (“yes” or “no”) with a right- or left-hand button press. Mapping between choice and response was instructed by an informative cue either before (pre-condition, cue 1) or after the stimulus (post-condition, cue 2). Additionally, there was an irrelevant cue offering no additional information either after (pre-condition, cue 2) or before (post-condition, cue 1). Participants additionally used the same button press to indicate their decision confidence, using an inner or an outer button. **(B)** Difference between relative reaction times depending on task, choice, stimulus, response and accuracy. For each comparison, all other variables were accounted for, and the difference in reaction times was computed after normalizing by the average of both options. Darker and brighter dots indicate participants performing task versions A and B, respectively. Horizontal and vertical bars indicate mean +/- SEM across participants.

### Decoding neural representations of stimulus, response and choice-response mapping

For each task condition separately, we quantified neural information about the stimulus, response, choice-response mapping and choice using a multivariate analysis approach (cross-validated MANOVA (Allefeld and Haynes, 2014; Christophel et al., 2018), Fig. S1). This method is a generalization of the commonly used cross-validated Mahalanobis distance. cvMANOVA builds on a multivariate general linear model to assess the cross-validated variability contained in the data that is related to a specific variable of interest. While conceptually similar to decoding algorithms, cvMANOVA offers a number of advantages. Firstly, it allows for the simultaneous extraction of information about multiple variables without repeatedly training decoders on each variable separately. Secondly, this enables the quantification of information related to one variable, while excluding confounds related to any other variable. Thirdly, the resulting measure of the separability of the multivariate activity patterns associated with the variables of interest is continuous, offering a better interpretability and higher sensitivity compared to classifier accuracy. In addition, cross-validation ensures the unbiased estimation of information by using non-overlapping test- and training data sets. Thus, importantly, this analysis isolated neural information about each individual variable, independently of the others. Choice information, for example, was the information contained in the neural data about a participant’s perceptual choice independent of all other variables.

We found significant neural information about all task variables in both conditions (P < 0.01, cluster-permutation statistics, Fig. 2). Stimulus information (i.e. the neural pattern distinctness between “signal” and “noise” trials) rose after stimulus onset and remained partially present after stimulus offset. Response information (i.e. right vs. left-hand button presses) built up after stimulus offset; it did so earlier in the pre-condition where the choice-response mapping was already known during stimulus presentation. Motor responses could be predicted more easily, and earlier in the trial, from motor-cortical beta lateralization (Fig. S2, (Pape and Siegel, 2016). Choice-response mapping information (i.e., yes/left and no/right vs. yes/right and no/left trials) peaked upon presentation of the relevant cue; after the pre-cue in the pre-condition and after the post-cue in the post-condition. Notably, in the pre-condition, mapping information was still present late in the trial, several seconds after presentation of the visual cue, indicating that mapping information was likely not purely sensory driven by the visual features of the cues.

**Fig. 2.**
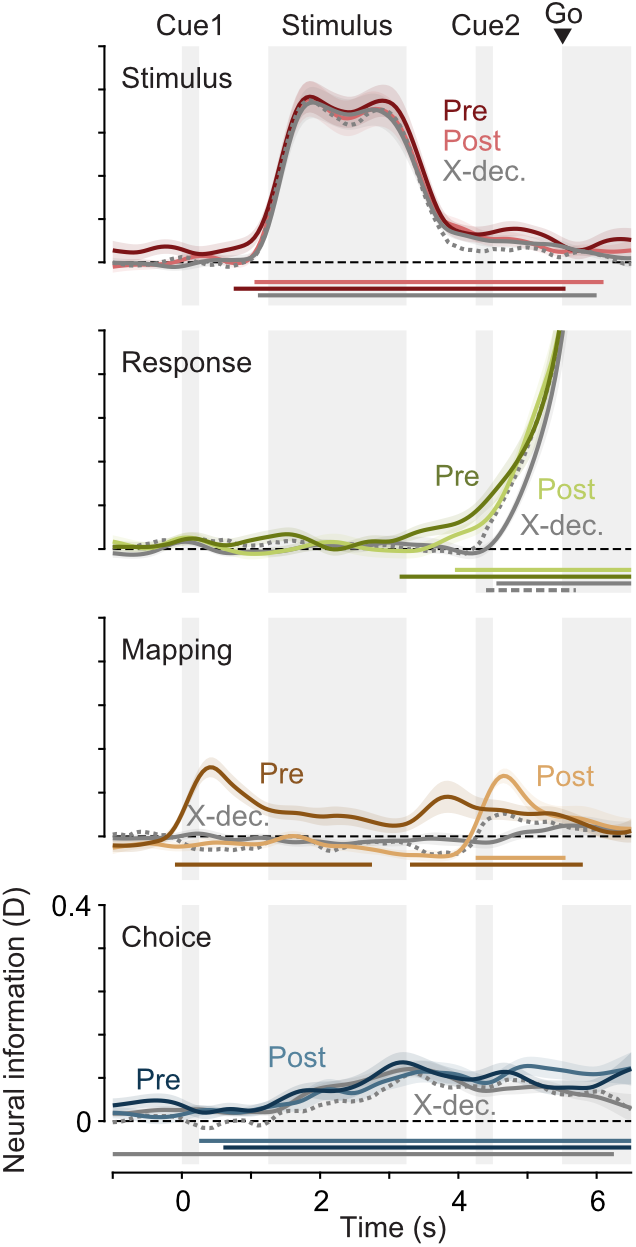
Neural information about the stimulus, response, mapping and choice. Darker lines indicate information during the pre-condition, brighter lines during the post-condition. Gray lines show the cross-decoding (‘X-dec.’) between both conditions, dashed gray lines the cross-decoding expected if representations in both contexts were identical. Horizontal lines denote temporal clusters of significant information (colored lines, P < 0.01, cluster-permutation, one-tailed, N = 26), cross-information (gray, two-tailed) or significantly less cross-information than expected (dashed gray, one-tailed). Coloured lines and shaded regions indicate the mean +/- SEM of information across participants.

### Abstract choice representations generalize between task contexts

Crucially, we also found information about the perceptual choice (i.e. yes vs. no choices, Fig. 2, bottom, *P* < 0.0001 in both ‘pre’ and ‘post’ conditions, cluster permutation). Even though participants’ choices were related to the presented stimuli and behavioral responses, our analysis framework ensured that choice information could not be explained by neural variability due to either stimuli or responses. Thus, choice information was stimulus- and response-independent. In both conditions, choices could be predicted before stimulus onset (pre: *P* = 0.003, post: *P* = 0.045; one-tailed t-tests on time-averaged choice information up to 1.25s), indicating that they were partly based on purely internal priors.

While, in the ‘post’-condition, the required motor action was not specified until after the stimulus, choices could be immediately mapped to the appropriate response in the ‘pre’-condition. Nevertheless, choice information was present in both conditions with a similar magnitude and time course (*P* > 0.05 for all time points before the end of the stimulus, two-tailed t-test), rising during stimulus presentation and remaining present until the end of the trial. Choice information could not be explained by eye movements (Fig. S3). Thus, choices were represented abstractly in the human brain, regardless of whether they could be directly linked to an action or not.

We employed a cross-decoding approach to assess the extent to which these choice representations were similar between both task conditions. We trained a decoding model on one task condition and tested it on the other. As the information estimated using cvMANOVA is symmetric with respect to the test- and training data used, we averaged results from both directions for all cross-decoding analyses. If the multivariate neural patterns distinguishing choices were identical in the ‘pre’- and ‘post’-conditions, we would expect the magnitude of the resulting cross-information to be comparable to the information found within the individual conditions. If, on the other hand, choices were represented in orthogonal neural subspaces in both conditions, cross-information should be much lower or negligible.

Cross-decoding of choices was positive throughout the trial (*P* < 0.0001, cluster permutation). Furthermore, the magnitude of cross-information was similar to the magnitude of choice information in the ‘pre’- and ‘post’-conditions. To quantify this, we derived an estimate of the expected cross-information under the assumption of identical representations in both conditions, i.e. representations relying on the same multivariate pattern and differing only in signal to noise ratio between conditions (see Materials and Methods). We found that cross-decoded choice information was never significantly lower than expected if representations were identical (*P* > 0.05 for all time points). Thus, abstract choice representations were not only present but were also shared between an action-linked and an action-independent choice context.

### Choice representations dynamically shift from sensory to motor areas

We further investigated the properties of neural stimulus, choice and response representations by pooling data from both task conditions. This choice was justified by our finding of shared choice representations and maximized the signal to noise ratio for the following analyses. We repeated the decoding analysis in a searchlight fashion across cortex to extract the spatiotemporal evolution of neural information about each variable (Fig. 3). During stimulus presentation, stimulus information was strongest in occipital visual cortex, in line with early visual representations of the sensory input. After stimulus offset, information remained at a lower level, uniformly across the brain (Fig. 3A, top). Response information increased earliest and most strongly in motor areas (Fig. 3A, middle), consistent with preparatory activity related to the upcoming motor response.

**Fig. 3.**
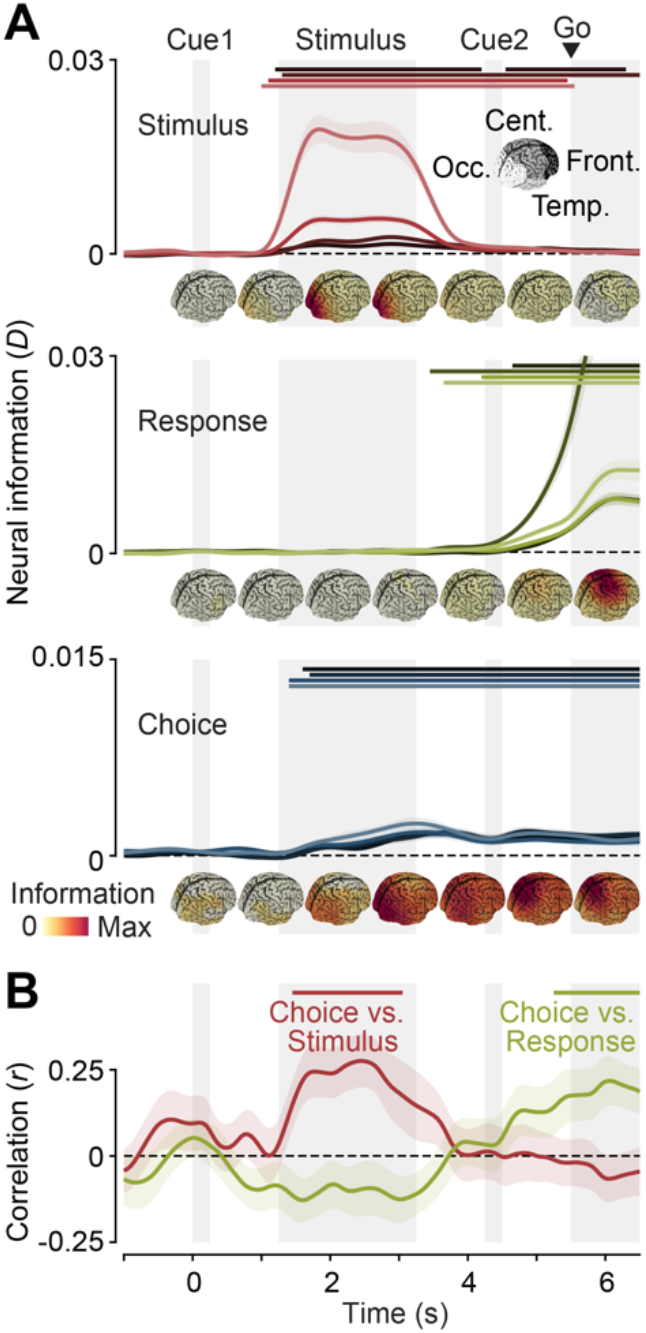
Spatiotemporal dynamics of neural information. **(A)** Time-resolved stimulus (top), response (middle) and choice (bottom) information in four groups of sources (in descending order of brightness: occipital, temporal, central, frontal). Data from both hemispheres was averaged. The cortical distribution of information during different time intervals is shown underneath the time-courses. **(B)** Correlation of the cortical distribution of choice information with the distribution of peak stimulus information (red) and peak response information (yellow). Horizontal lines denote temporal clusters of significant information (A, *P* < 0.05, cluster-permutation, one-tailed, N = 26) or correlation (B, *P* < 0.05, cluster-permutation, one-tailed, N = 26). Coloured lines and shaded regions indicate the mean +/- SEM of information or correlation across participants.

The expected cortical distribution and temporal evolution of choice information is less clear. Choices may be represented in visual areas, consistent with findings of choice probabilities in sensory neurons reflecting either the effect of sensory noise on decision formation or high-level feedback onto sensory populations (Britten et al., 1996; Nienborg and Cumming, 2009; Siegel et al., 2015). Choice-specific signals may also be present in motor- and premotor areas, supporting the planning of potential motor responses (Bennur and Gold, 2011; Cisek and Kalaska, 2005; Herding et al., 2017; Siegel et al., 2015) or in associative areas specialized for decision formation.

We found that the distribution of choice information changed dynamically over the course of the trial, rising first in occipital areas, before spreading throughout the brain. After the go cue, choice information remained strongest in parietal cortex and central motor areas (Fig. 3A, bottom). Given the apparent shift of choice information from occipital areas during stimulus presentation to central areas during the response phase, we quantified the similarity the cortical distribution of choice information exhibited with those of stimulus and response information. We found a significant correlation between the cortical distributions of choice and stimulus information during stimulus presentation, and between choice and response information during the response phase (Fig. 3B, stimulus: *P* = 0.0064, response: *P* = 0.0227, cluster permutation). We found similar results when repeating the searchlight analysis independently for pre- and post-condition trials and extracting correlation values for the early stimulus-related and the later response-related cluster. Despite the reduced number of trials, two out of four correlation values were significant, and all four had the same directionality as in the pooled data (stimulus vs. choice in pre: *t_25_* = 4.24, *P* = 0.0001, response vs. choice in post: *t_25_* = 2.81, *P* = 0.0047, stimulus vs. choice in post: *t_25_* = 0.98, *P* = 0.1683, response vs. choice in pre: *t_25_* = 1.64, *P* = 0.0569, all one-tailed t-tests).

### Temporal stability of neural representations

The spatial overlap between choice, stimulus and response information raised the question whether there were shared representations between stimulus and choice during evidence accumulation and between choice and response during motor execution, respectively. We used cross-temporal and cross-variable decoding to test this and evaluated both the temporal dynamics of representations and the relationships between stimulus, choice and response representations (Fig. 4A).

**Fig. 4.**
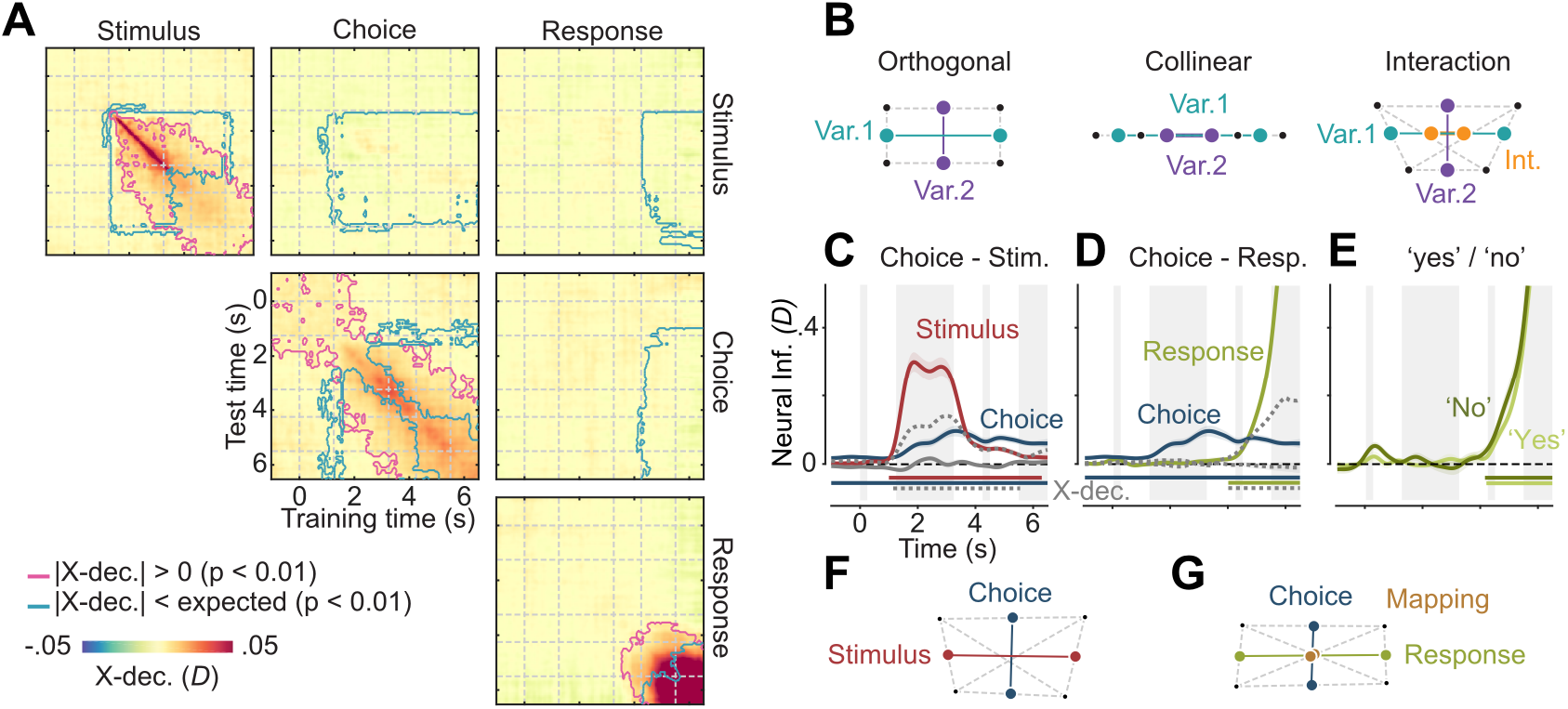
Relationship between stimulus, choice, and response representations. **(A)** Cross-temporal and cross-variable decoding. Colors indicate neural information when trained and tested on any pair of time-points and variables. Pink outlines indicate clusters of shared information between timepoints and variables, i.e. pairs of time-points and/or variables during which cross-information is significantly different from 0 (|X-dec.| > 0, cluster-permutation, *P* < 0.01, N = 26, two-tailed), blue outlines indicate different representations between timepoints and variables, i.e. pairs of time-points and/or variables during which cross-information is significantly smaller than expected for identical representations (|X-dec.| < expected, one-tailed). **(B)** Possible relationships between the representations of two variables. Points indicate average activity patterns for different conditions, distances between points the strength of information. Representations may be orthogonal, collinear, or orthogonal but linked with an interaction. **(C)** and **(D)**, Cross-variable decoding between choice and stimulus, and choice and response, respectively. Colored lines show neural information about each variable, gray lines cross-variable information (X-dec.), and dashed gray lines the expected cross-information if both variables were represented identically. Horizontal lines indicate clusters of significant information (colored, *P* < 0.01, one-tailed), or significantly less cross-information than expected (dashed gray, *P* < 0.01, one-tailed). **(E)**, Response information for “yes” and “no” choices. Coloured lines and shaded regions in panels C, D and E indicate the mean +/- SEM of information across participants. **(F)**, Visualization of the relationship between stimulus and choice representations, based on the cross-decoding values in (C). Stimulus and choice are nearly orthogonal. **(G)**, Visualization of the relationship between choice and response representations, including mapping as their interaction, based on (D) and (E). Choice and response representations are nearly orthogonal, and response representations are equally strong for both choices. Thus, there is no systematic relation between the neural patterns encoding choice, stimulus and response.

First, we focused on the temporal dynamics of representations. By using data from one timepoint for training, and from another timepoint for testing, cross-temporal decoding can reveal time periods of relative stability (King and Dehaene, 2014). Furthermore, it is possible to compute the expected cross-temporal decoding under the assumption that the underlying representation remains perfectly stable over time. Comparing the empirical cross-temporal decoding to this expectation can reveal periods of relative dynamics (Spaak et al., 2017). Stimulus information was initially highly dynamic, as indicated by high cross-decoding values being concentrated along the diagonal, but stable after stimulus offset, as indicated by significant cross-decoding far from the diagonal (Fig. 4A, top left). Given our use of fixed random dot patterns, this was consistent with stimulus information being driven by two components: During stimulus presentation, information was likely dominated by moment-to-moment differences in retinal input. After stimulus offset, the global motion content may have contributed more strongly. Choice information was temporally more stable; still, early and late choice representations were distinct (Fig. 4A, center), in line with the observed spatial shift from sensory to motor areas.

### Choice representations are distinct from sensory and motor representations

How did the neural representations of different variables relate to each other? The multivariate patterns that encode any two variables are either orthogonal, indicating non-overlapping underlying population subspaces, collinear, indicating indistinguishable circuits underlying both representations, or somewhere in between (Fig. 4B; see also Fig. S1 for further details). Furthermore, the representation of one variable may differ depending on the value of the other, i.e. the two variables may interact. In the present data, stimulus- and choice representations may depend on identical underlying circuits. For example, sensory neurons may show the same responses for visually presented as for imagined motion (Zhao et al., 2020; Zhou and Freedman, 2019). If such neurons constituted stimulus- and choice representations, we would expect strong positive cross-information between stimulus and choice. In contrast, if choice and stimulus information were largely driven by distinct populations, this may result in no cross-information; Our results were compatible with the latter scenario. There was no significant cross-decoding between stimulus and choice (Fig. 4A, top center, biggest cluster: *P* = 0.11; Fig. 4C, biggest cluster: *P* = 0.22), and cross-decoding was significantly lower than expected for identical representations (Fig. 4A, top center; Fig. 4C, *P* < 0.0001).

Next, we investigated the relationship between choice- and response representations. Again, we found only weak cross-information between the two variables, indicating that neural choice and response representations did not overlap (Fig. 4A, middle right, biggest cluster: *P* = 0.15; Fig. 4D, biggest cluster: from 0.8 to 2.55 s, *P* = 0.038, uncorrected). After selection of a motor response, choices may still have been represented as a modulation of the motor signal, e.g. leading to a relative strengthening of the activity pattern associated with the upcoming motor response for “yes”-choices compared to “no”-choices. We thus assessed the magnitude of response information, separately for each choice. However, we found no difference between both conditions (*P* > 0.05 for all time points), indicating that even during response execution, choices were not represented as a modulation of neural motor activity. (Fig. 4E). We further visualized these results geometrically, which well illustrated the near-orthogonality of choice- and stimulus-, or choice- and response signals, respectively (Fig. 4F-G). In sum, the neural circuit patterns underlying choice information in our MEG data were not significantly shared with those underlying stimulus and response information, even when they were strongest in similar areas.

Abstract choice signals may also be related to, and caused by, sequential choice biases, i.e. preceding choices (Lueckmann et al., 2018; Siegel et al., 2015; Urai et al., 2019). Furthermore, when pooling over the pre- and post-conditions, the higher signal-to-noise ratio revealed robust pre-stimulus choice information (Fig. 4A, C, D), indicating the formation of choices even before stimulus presentation, which may well be related to sequential choice biases. We therefore repeated the analysis including the previous choice as an additional variable, and found that choice information after stimulus onset could not be explained by the previous choice (Fig. S4A). Including the previous motor response instead of previous choice showed a sustained representation of past motor actions (Pape and Siegel, 2016). However, this had an even weaker effect on choice information. Thus, neuronal choice signals did not merely reflect the previous choice or motor response.

While our analysis already excluded the possibility that choice information was driven by overall differences between stimuli, it could theoretically still be explained by a difference between correct and error trials for one of the two stimulus classes. To eliminate this possibility, we trained the choice decoding model on all trials and evaluated it separately on correct and incorrect trials. As choice information was present in both cases, and had the same sign, it could not be explained by choice accuracy (Fig. 5). In sum, abstract choice information did not result from the representation of either previous choices or accuracy as potentially confounding variables.

**Fig. 5.**
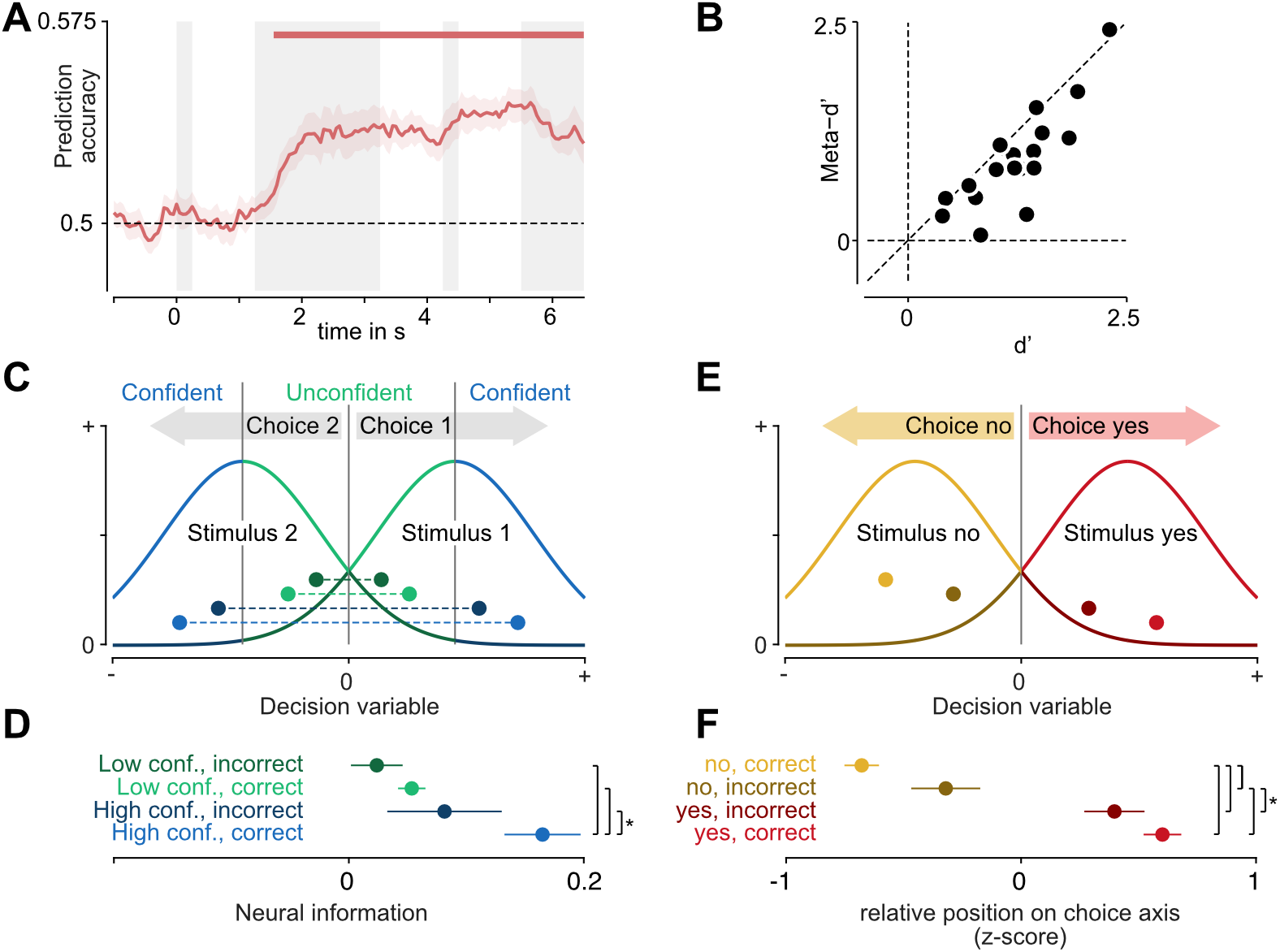
Choice representations behave like a decision variable. **(A)**, Prediction of stimulus class from the sign of single trial choice information. Mean +/- SEM across participants **(B)**, Behavioral sensitivity (d’) and meta-cognitive sensitivity (meta-d’). **(C)**, The relationship between decision variable, confidence, and accuracy as predicted by signal detection theory. For each of the two stimuli, the distribution of values of the decision variable is centered on the respective side of the decision boundary at 0. When the absolute distance to the decision boundary is larger, the observer is more confident in their choice. Correct and incorrect, confident and unconfident trials are color-coded as in (D). **(D)**, Time-averaged choice information (1.25 to 4s) in trials split by confidence and accuracy. Stars denote significance (*P* < 0.05, one-tailed t-test, N=19). **(E)**, The relationship between decision variable, accuracy and choice as predicted by signal detection theory. For both yes- and no-choices, the decision variable has a higher absolute magnitude in correct trials. Correct and incorrect, yes and no trials are color-coded as in (F). **(F)**, Time-averaged (1.25 to 4s), normalized placement on the choice axis of trials split by choice and accuracy. Stars denote significance (*P* < 0.05, one-tailed t-test, N=19).

### Choice signals predict stimuli

The above results suggested that choice signals reflected an abstract decision stage which was neither directly related to early sensory nor to motor representations. Nonetheless, behavioral choices were based on the presented stimuli, and therefore strongly correlated with them. We thus asked whether this relationship was reflected in the neural choice signals, or whether they were stimulus-independent and therefore purely internally driven. To do so, we computed single-trial estimates of the choice signal by projecting each trial’s data onto the choice axis defined by our multivariate analysis. We then assessed whether the sign of the single-trial choice signals predicted the stimulus. Indeed, after stimulus onset, the predictability of the stimulus increased until it reached a stable level for the remainder of the trial (Fig. 5A, P < 0.0001, cluster-permutation). As expected, there was no significant stimulus-predictability before stimulus onset, even though there was a small amount of choice information (Fig. 4C).

### The strength of choice signals predicts decision confidence and accuracy

Our participants also reported their confidence in each trial’s perceptual choice, providing us with further leverage to unravel the nature of the choice signals we found. Specifically, this allowed us to test whether the choice-predictive signal merely correlated with choices, or whether its relation to accuracy and confidence exhibited additional key properties expected of a decision variable integrating evidence towards a choice.

First, we behaviorally assessed the relationship between participants’ choices and confidence ratings. Participants were more confident in yes-than in no-choices (average proportion of high confidence reports: 0.54 vs. 0.47, *P* = 0.034, two-tailed t-test) and in trials with signal-than in those with noise stimuli (0.55 vs. 0.46, *P* = 1.4×10^−4^). In addition, and critically, they reported high confidence more often in correct trials than in incorrect trials (average proportion of high confidence reports: 0.56 vs. 0.35, *P* = 3 ×10^−7^, two-tailed t-test). We quantified this relationship using the meta-d’ measure of metacognitive sensitivity (Maniscalco and Lau, 2012) (Fig. 5A). As expected, meta-d’ was positive (*t_18_* = 7.4, *P* = 7.5×10^−7^, two-tailed t-test), correlated with d’ (*r_17_* = 0.82, *P* = 1.5×10^−5^, Pearson correlation), but tended to be smaller than d’ (*t_18_* = −4.1, *P* = 7×10^−4^, two-tailed t-test). This showed that participants veridically reported their confidence and suggested that their confidence judgements were largely, but not perfectly based on the same sensory evidence as their choices (Grimaldi et al., 2015; Kiani et al., 2014; Maniscalco et al., 2021; Odegaard et al., 2018). These results also held when we separately assessed them in the pre- and post-conditions, and neither d’ (*t_18_* = 1.2, *P* = 0.24) nor meta-d’ (*t_18_* = 0.2, *P* = 0.83) were significantly different between conditions.

Next, we directly investigated the relation of neural choice signals to decision confidence and accuracy. In signal detection theory and in related accumulator models of decision making an internal decision variable tracks the integrated evidence for a given choice. Importantly, such a decision variable enables the computation of choice-confidence, as the absolute distance to the decision boundary (Hebart et al., 2016; Kiani and Shadlen, 2009; Macmillan and Creelman, 2004; Shadlen and Kiani, 2013). Consequentially, the absolute value of the decision variable should be larger during high-confidence trials than during low-confidence trials (Fig. 5B, blue vs. green), and, importantly independent of confidence, higher during correct than during error trials (Fig. 5B, bright vs. dark colors).

To establish whether the choice signals found here could reflect an internal decision variable, we thus repeated our decoding analysis, now adding decision confidence as an additional variable. We trained the decoding model separately on confident and unconfident trials, and tested it separately on confident correct, unconfident correct, confident error and unconfident error trials. We hypothesized that, if choice information constituted an internal decision variable reflecting the same subjective evidence used to inform confidence judgements, it would be strongest in correct trials when confidence was high, and progressively weaker in confident error trials, unconfident correct trials and unconfident error trials.

Indeed, we found that the strength of choice representations descriptively followed this pattern predicted by signal detection theory (Fig. 5C: correct/high confidence larger than incorrect/high confidence, correct/low confidence and incorrect/low confidence; *P* = 0.013, *P* = 0.002, *P* = 0.001; high confidence larger than low confidence and correct larger than incorrect; P = 0.027, P = 0.025). Importantly, participants’ accuracy and confidence were assessed as separate factors. Thus, the relationship between choice and confidence could not simply be explained by accuracy or vice versa. We additionally performed this analysis separately for the pre-cue and post-cue task conditions, after excluding the factor of response from our model in order to retain a sufficient number of trials per condition. There was a similar pattern in both tasks (Post: correct/high confidence larger than incorrect/high confidence, correct/low confidence and incorrect/low confidence; *P* = 0.035, *P* = 0.008, *P* = 0.005. Pre: correct/high confidence larger than incorrect/high confidence, correct/low confidence and incorrect/low confidence; *P* = 0.007, *P* = 0.002, *P* = 0.023). In contrast, there was no clear relationship between choice confidence or accuracy and the strength of stimulus- or motor representations (Fig. S5).

Finally, we investigated the relative placement of correct and incorrect trials of both choices on the neural choice axis. As the stimulus design used was inherently asymmetric (signal vs. noise stimuli), a similar asymmetry may be expected for the neural representations of yes- and no-choices, opening the door for potential, choice-unrelated confounds. For example, the timing of choice commitment may be different for yes- and no-choices, differentially affecting the neural signal. While our fixed-time design did not provide access to commitment times, responses in similar forced-response detection tasks have been found to be slower for no-than for yes-choices, and highly similar between correct and incorrect no-choices (de Gee et al., 2014). This is consistent with no-choices occurring when the internal decision variable does not hit a bound until the end of the trial. This leads to a critical prediction for the present data. If neural choice signals reflected the time of choice commitment rather than the decision variable itself, they should exhibit a similar pattern with a difference between yes- and no-choices but similar signals for correct and incorrect no-choices. To test this, we trained a decoding model on all choices and tested it separately on, first, correct yes vs. correct no-choices, secondly, incorrect yes vs. incorrect no-choices, and third, correct yes vs, incorrect no-choices. The resulting distances allowed us to estimate the relative placement of correct and incorrect, yes and no choices on the choice axis. As expected from a neural decision variable, these trial types were well-ordered, with correct no-choices being followed by incorrect no-, incorrect yes-, and correct yes-choices (Fig. 5E-F, P<0.05 for all pairwise comparisons apart from correct yes vs. correct no with P = 0.12, one-tailed t-tests). In contrast, this ordering is incompatible with a timing confound arising from a potential asymmetry between yes- and no-choices.

In sum, our behavioral results pointed to the existence of an internal decision variable, which informed both choices and confidence ratings. Furthermore, the strength of the choice-predictive neural signal varied with confidence and accuracy, precisely following a pattern predicted from signal detection theory. Thus, neural choice information measured in MEG did not only predict abstract perceptual choices but appeared to reflect the underlying internal decision variable.

## Discussion

Studies of the neural basis of sensorimotor decision-making have often neglected abstract, motor-independent choices (Donner et al., 2009). This is rooted in the fact that many real-world choices appear to be choices between motor actions (Cisek and Kalaska, 2010) and in the difficulty of accessing signals representing purely abstract choices. On the one hand, in animal studies, which provide the majority of evidence in support of neural circuits selective for specific choice options, behavioral tasks that disentangle choices from motor responses are very challenging. Non-invasive human studies, on the other hand, struggle to read out choice contents, and thus mostly provide indirect evidence for choice-related neural activity. Consequently, studies comparing the representation of choices in abstract and action-linked contexts are rare. A small number of notable exceptions have provided intriguing results (Bennur and Gold, 2011; Ludwig et al., 2018; Wang et al., 2019), but not established a unified account of the role and extent of abstract choice signals.

By combining non-invasive MEG in humans with an advanced multivariate analysis framework, we robustly read out abstract choice contents from whole-brain neural activity. In accordance with current theories (Kiani and Shadlen, 2009; Shadlen and Kiani, 2013), abstract choice representations predicted decision confidence and accuracy. This indicates that this neural signal did not merely correlate with choice but reflected the underlying decision variable. In sum, our findings point to an important role of abstraction in decision-making, even in a simple task involving a known sensorimotor mapping.

Abstract choices were not only represented in brain activity when decisions had to be made abstractly, but also when the sensorimotor mapping was known in advance. Importantly, our cross-decoding analysis showed that choice representations in both task contexts were indistinguishable from one another. While the fundamental limits of MEG spatial resolution and sensitivity prevent the conclusion that the underlying circuit representations are identical, this striking similarity requires any potentially remaining differences between conditions to be small and of a type that MEG is blind to.

Our finding of abstract choice representations generalizing between contexts in which actions can be planned and those in which they cannot is in line with behavioral evidence suggesting analogous mechanisms underlying decision-making in action-linked and -independent contexts (Tsetsos et al., 2015). Moreover, recordings in macaque area LIP have found the choice selectivity of neurons to be similar, regardless of whether a motor action was specified (Bennur and Gold, 2011). Our results extend this finding to the whole-brain level, indicating that the dominant sources of choice-selective signals generalize between contexts. Intriguingly, a recent study found representations of the decision variable in area LIP that were not tightly linked to the population’s oculomotor selectivity, but varied in a task-dependent manner (Okazawa et al., 2021). These task-dependent representations are compatible with abstract, motor-independent choice representations as reported here. Furthermore, our findings accord well with research implicating a centro-parietal positivity (CPP) as an electrophysiological marker of evidence accumulation (O’Connell et al., 2012; Twomey et al., 2016). The CPP exhibits several properties of a domain-general, abstract neural decision variable; however, while it gradually builds up with the absolute amount of evidence, it has not been shown to carry information about the choice itself. Thus, the CPP, as an unsigned marker of integration, and the specific choice signals found in the present study may reflect different aspects of the same underlying process.

This - as well as any other - decision making study lives off the fact that sometimes participants make different choices for identical stimuli. How does this variability arise? In principle two broad, and not mutually exclusive, classes of explanations exist. First, it could be bottom-up driven, with sensory noise having a causal effect on choices. This sensory noise could be internal, arising from the inherent variability in neural responses for identical stimuli, or due to uncontrolled external variability, such as small differences in the stimulus itself. Secondly, it could be top-down driven, with internal factors such as expectations, biases or beliefs or simply non-sensory noise pushing choices one or the other way. Several of our results consistently suggest that the demonstrated choice signals are positioned at an intermediate stage between these extremes. First, if the choice signals directly reflected sensory noise, we would expect this noise to inhabit the same neural subspace as the stimulus signals themselves - in other words, there should be strong cross-information between stimulus and choice. In contrast to this, our results are better compatible with choice signals reflecting integrated sensory noise represented distinctly. For example, one may expect to find instantaneous sensory noise represented in area MT, but integrated sensory information, and therefore integrated noise as well, represented in area LIP. Notably, such an integration stage would still be expected to be modulated by the stimulus, and thus to lead to stimulus-choice cross-information, but only subtly so. In our data this effect was too weak to result in significant cross-information, but is nonetheless apparent in the hypothesis-driven finding of stronger choice signals in correct than in error trials. Second, we found that signed choice information predicted the stimulus - despite near-orthogonality of the representations of both variables. This suggests that choice information was indeed reflective of a stage separate from, but influenced by the early sensory representation. This prediction increased during stimulus presentation and then remained stable, similar to the choice information time course itself, and consistent with the time course expected from temporal integration. In contrast, choice signals at an instantaneous, early sensory non-integration stage would also predict the stimulus, but predictions should be at the same level throughout the stimulus presentation interval, and subsequently taper off. Third, we found small amounts of choice information before stimulus presentation. As these could not have arisen due to the stimulus, they must reflect intraneous factors. In conclusion, the most parsimonious explanation for our data is an intermediate choice stage which reflects both accumulated sensory evidence and top-down contributions, akin to an internal decision variable.

The stimuli used in the present study, and therefore the corresponding choices, were inherently asymmetric. Thus, one may plausibly assume that this asymmetry underlies the decodablity of choices: an unobserved, confounding variable correlated with choice may result in the seeming readout of choices. However, our finding of a significant difference of the choice signal between correct and incorrect no-choices does not accord with a timing-related confound due to this asymmetry (de Gee et al., 2014). More generally, any confounding variable would have to exhibit the properties of the choice signal demonstrated here: small, but existing pre-stimulus differences between choices, a modulation by confidence and accuracy even within no-trials, and a trial-by-trial predictability of the stimulus. We therefore consider it unlikely that our results can be accounted for by the asymmetric task-design. Nonetheless, future research should explicitly test whether the current findings generalize to symmetric choices.

The cortical distribution of abstract choice signals may be modulated by response modality. Recent work using fMRI suggested that, for vibrotactile comparisons, abstract choice representations are present in non-overlapping, modality-specific cortical areas (Wu et al., 2019, 2021). On the other hand, direct neuronal recordings have shown the representation of recognition and categorization choices in medial frontal cortex to generalize between manual and saccadic responses (Minxha et al., 2020). The accessibility of such modality-independent representations of choice likely depends on the specific behavioral task and type of neural measurement. Research combining multiple measurement scales (Cichy et al., 2014; Kriegeskorte et al., 2008; Sandhaeger et al., 2019) should help resolving this. Our results only have indirect implications for the modality-dependence of choice signals because participants eventually always reported their choice using a button press. Nonetheless, the broad availability of choice representations across the brain, in conjunction with the shift of the information peak from visual sensory to motor areas is consistent with the co-existence of modality-independent and modality-specific components.

Perceptual decisions involve a complex interaction of feedforward and feedback processes throughout the brain (Quinn et al., 2021; Siegel et al., 2015; Wilming et al., 2020). Here, we found that the spatial peak of abstract choice information shifted throughout the trial, reflecting the currently relevant stage (Li Hegner et al., 2017; Siegel et al., 2015). This does not necessitate that choices originate in sensory cortex and are later relayed to motor cortex; in fact, choices may be computed elsewhere, but be preferentially accessible in currently engaged areas. The global availability of choice information is in line with either a distributed computation that involves recurrent interactions, or a global broadcast of choice signals (Nienborg and Cumming, 2009; Siegel et al., 2015), for example through feature-attentional mechanisms (Quinn et al., 2021). Further studies including invasive and manipulative approaches are required to pinpoint where and by which mechanisms abstract choices are computed.

A growing body of evidence has related the formation of action-linked sensorimotor decisions to activity in motor- and premotor areas (Cisek and Kalaska, 2005; Donner et al., 2009; Gold and Shadlen, 2000; Shadlen and Newsome, 2001; Wang et al., 2019; Wilming et al., 2020). Our findings are well compatible with these results: the presence of an abstract choice stage does not preclude the simultaneous planning and competition of multiple response options or a general unspecific response preparation (Cisek and Kalaska, 2005; Klaes et al., 2011). Indeed, we found fluctuations in motor cortical beta band activity to predict upcoming motor responses, independently of the perceptual choice (Pape and Siegel, 2016; Pape et al., 2017). These response-predictive beta band signals ramped up upon stimulus presentation in the “pre”-condition, as expected due to the earlier availability of the choice-response mapping. Notably, this ramp-up happened earlier than the appearance of response information in the broadband electrophysiological signals, underpinning the well-known role of beta band activity as a specific marker of motor preparation (Twomey et al., 2016). Taken together, these findings support a multi-level model of decision-making involving simultaneous evaluation of abstract choices as well as motor actions (Cisek, 2012). The relevance of an abstract choice level may be understood in light of phenomena like perceptual priors (Haefner et al., 2016; Summerfield and de Lange, 2014), sequential choice biases (Pape and Siegel, 2016; Urai et al., 2019), or value computations associated with the choices themselves (Padoa-Schioppa, 2011), which all require and act on abstract choice representations. The primate brain, which is able to assess abstract options and treat decision-making problems as arbitrary categorization (Freedman and Assad, 2011; Merten and Nieder, 2012), may do so even when not strictly necessary. Importantly, this can still be reconciled with an intentional framework of decision-making, if intentions are not only about actions, but also rules, or activations of neural circuits in general (Cisek, 2012; Shadlen and Kiani, 2013).

We conclude that an abstract choice stage may be universally present in human perceptual decision-making, enabling the evaluation of motor-independent choice options even during action-linked decisions.

## Methods

### Experimental Design

#### Participants

33 healthy, right-handed human volunteers (18 female; mean age: 28 years; 3 years SD) participated in this study and received monetary reward. All participants had normal or corrected-to-normal vision and gave written informed consent before participating. The study was conducted in accordance with the Declaration of Helsinki and approved by the ethics committee of the University of Tübingen.

#### Behavioural task & stimuli

Participants performed a flexible sensorimotor decision-making task. In each trial, they had to decide whether a random dot kinematogram contained coherent downwards motion or not and reported their choice with a left- or righthand button press. Crucially, the mapping between response hand and choice varied on a trial by trial basis. Moreover, the mapping was either revealed before (pre-condition) or after (post-condition) the stimulus. Additionally, an irrelevant cue was presented after (pre-condition) or before (post-condition) the stimulus.

Participants started a trial by acquiring fixation on a fixation spot. After a fixation period, the first cue appeared for 250 ms, followed by a delay of 1000 ms, the presentation of the random dot stimulus for 2000 ms, another 1000 ms delay and the second cue for 250 ms. A third 1000 ms delay was followed by a 33 ms dimming of the fixation spot, which served as the go-cue for the participant’s response. The response consisted in a button press using the left or right index finger, according to the choice and the choice-response mapping. Participants chose one of two buttons on either side to indicate whether they were confident in their choice or not. 250 ms after their response, participants received a 100 ms visual feedback (centrally presented circle, 2.1 degree diameter, red for incorrect or green for correct).

The random dot stimuli consisted of 1500 white dots with a diameter of 0.12 degrees, presented in an 8.5 degree diameter circular aperture on a black background. Dots moved at a speed of 10 degrees per second. For each participant, we used only two stimuli, each presented in half of the trials: First, a target stimulus, in which, on each frame, a fraction of dots moved coherently downwards, whereas the rest moved in random directions. Secondly, a noise stimulus, in which all dots moved in random directions. In a separate session before the MEG recordings, the motion coherence of target stimuli was titrated to each participant’s individual perceptual threshold using a staircase procedure with 280 trials. Motion coherence was adaptively lowered by one level after each correct choice and increased by two levels after each incorrect choice. To determine the coherence threshold, a Weibull function was fit to the resulting data, excluding the first 50 trials. Choice-response cues and irrelevant cues all had the same luminance and size (0.85 degree diameter).

Each participant took part in two recording runs of one of two task versions which differed in the details of the choice-response cue as well as the confidence report. Participants 1-20 performed version A: here, the choice-response cue consisted of a centrally presented red or green square (yes=right hand: green; yes=left hand: red), whereas the irrelevant cue was a blue square. The outer button always indicated a confident, the inner one an unconfident choice. In this version, the fixation baseline at the beginning of each trial lasted 1500 ms. Each recording run consisted of 400 randomly ordered trials, of which 120 were pre-cue trials, 120 post-cue trials, and 160 belonged to one of two control conditions not reported here. Participants 21-33 performed version B: here, the choice-response cue consisted of two vertical rectangles (yes=right hand: left rectangle mint, right rectangle pink; yes=left hand: left rectangle pink, right rectangle mint) forming a square, whereas the irrelevant cue consisted of two horizontal rectangles (upper: pink, lower: mint). The confidence mapping (inner or outer button for confident / unconfident responses) was changed in each recording run. Here, the fixation baseline was 1000 ms. Each run consisted of 400 randomly ordered trials, 200 of which were pre-cue and 200 post-cue trials. The changes in version B were designed to minimize sensory- and motor confounds in a separate analysis of task- and confidence-related effects (not reported here). The data from version A was previously used in another publication (Pape and Siegel, 2016).

To ensure that participants were performing both task conditions well, we computed overall accuracy as the percentage of correct trials. We used a two-tailed paired t-test to test whether accuracy was different between task conditions. To make sure participants did not systematically associate one of the motor responses with one of the choices, we computed the percentage of “right” button presses for “yes”- and “no” choices separately and compared both against 50% using two-tailed t-tests.

#### Setup & recording

We recorded MEG (Omega 2000, CTF Systems, Inc., Port Coquitlam, Canada) with 275 channels at a sampling rate of 2,343.75 Hz in a magnetically shielded chamber. Participants sat upright in a dark room, while stimuli were projected onto a screen at a viewing distance of 55 cm using an LCD projector (Sanyo PLC-XP41, Moriguchi, Japan) at 60 Hz refresh rate. Stimuli were constructed offline and presented using the Presentation software (NeuroBehavioral Systems, Albany, CA, USA). To ensure continuous fixation, we recorded eye movements using an Eyelink 1000 system (SR Research, Ottawa, Ontario, Canada).

### MEG preprocessing and source analysis

#### Preprocessing

We low-pass-filtered MEG and eye-tracking data at 10 Hz (two-pass forward-reverse Butterworth filter, order 4) and down-sampled to 20Hz. Trials containing eye-blinks were rejected. We chose not to apply a high-pass filter in order to avoid filter artefacts (van Driel et al., 2021). At the same time, we could not use a baseline correction as choice effects could plausibly be driven by previous trials. We thus used robust detrending (de Cheveigné and Arzounian, 2018) to remove polynomial trends from the MEG data, but not the eye tracking data, in a piecewise fashion (600 s pieces, removal of linear trend followed by 10th order polynomial). Data of three participants was rejected due to metal artifacts.

#### Source reconstruction

For source reconstruction based on each participant’s individual anatomy, we recorded structural T1-weighted MRIs (echo time (TE) = 2.18ms, repetition time (TR) = 2.3 ms, longitudinal relaxation time (T1) = 1.1 ms, flip angle = 9°, 192 slices, voxel size 1×1×1 mm3) with a Siemens 3T Tim Trio scanner and a 32 channel Head Coil. We generated single-shell head models (Nolte, 2003) and estimated three-dimensional (x, y and z-direction) MEG source activity at 457 equally spaced locations 7 mm beneath the skull, using linear spatial filtering (Van Veen et al., 1997). We retained, for each source, activity in all three directions and concatenated the data of the two separate recording runs per participant. For all subsequent analyses, we reduced the dimensionality of this 1371-dimensional source space: for all whole-head decoding analyses we performed principal component analysis, retaining the 75 components with the largest variance across all combinations of task variables. For searchlight analyses, we used each of the 457 sources’ immediate neighbors, including all 3 dipole directions.

### Multivariate decoding analyses

#### Task variables and cross-validation scheme

The experimental design resulted in a number of variables of which each trial instantiated a combination. For each trial, we defined the task (pre- or post-cue), stimulus (target or noise), response (left or right hand button-press), mapping (target = left or target = right), choice (yes/target or no/noise), accuracy (correct or incorrect) and confidence (high or low). Not all of these variables were independent of each other: for a given stimulus and choice, accuracy is fixed; and for a given choice and mapping, response is fixed. Thus, 5 independent variables giving rise to 32 conditions remained (Fig. S1A). While those variables under experimental control (task, stimulus, mapping) were fully balanced, those dependent on the participants’ behavior (choice, response, confidence, accuracy) were not, leading to a non-uniform sampling of conditions (Fig. S1A). To ensure an accurate estimation of neural information about each variable, independent of the others, we implemented an n-fold cross-validation scheme, where n was the lowest trial count per condition. Thus, for each cross-validation fold, both training and test data contained trials of all conditions. In order to decrease the dependence of our results on a particular random partition into folds, we repeated each analysis 10 times, with different random seeds. All results were averaged across these random seeds before further processing.

Due to the variability in behavioral responses, as well as the rejection of trials containing eyeblink artefacts, we did not retain the same amount of trials from each condition for all participants. However, to accurately estimate neural information we needed to ensure that, first, there were trials of each condition, and second, the total number of trials was large enough in comparison to the dimensionality of the data to enable an unbiased estimate (Allefeld and Haynes, 2014). Specifically, each analysis requires at least N + K + 1 trials, where N is the number of channels and K is the number of independent variables in the model. For our main analyses (Fig. 2, Fig. 3, Fig. 4, Fig. S3, Fig. S3), including task, stimulus, choice, response, mapping and accuracy as variables, data from 26 participants had sufficient trials. When additionally including confidence as a variable, but neglecting the task condition (Fig. 5, Fig. S5), we retained 19 participants. To assess the effect of confidence separately for both task conditions, we used all variables apart from response, leading again to 19 useable participants. To assess the effect of the previous choice in relation to the current choice (Fig. S4A), we neglected the task condition as well as confidence, and included stimulus, choice, response, mapping, accuracy, and previous choice. This left us with data from 23 participants. To assess the effect of the previous motor response in relation to the current choice (Fig. S4B), we neglected the task condition as well as confidence, and included stimulus, choice, response, mapping, accuracy, and previous response. This left us with data from 25 participants. For all decoding analyses, we combined source level data from both recording runs per participant. Using source-level data allowed us to reduce between-run variance and reduce non-neural variability. To do so, we normalized the data per channel, time point and run over trials, and then concatenated data of both runs.

#### Cross-validated MANOVA

We used cross-validated MANOVA (Allefeld and Haynes, 2014; Christophel et al., 2018) to estimate the amount of information in multivariate MEG data about the task variables of interest. CVMANOVA estimates the variability explained by the task variables in relation to unexplained noise variability. Here, we re-implemented cvMANOVA for time-resolved data, adding the capability of cross-decoding by training and testing the model on different time points, variables, or levels of any variable. To this end, we first estimated a baseline noise covariance matrix, using trials from all unique conditions. We then “trained” the model by estimating contrasts of beta weights of each unique condition in a cross-validation fold’s training set, and “tested” it by estimating contrasts of beta weights in the fold’s test set. An estimate of true pattern distinctness was computed as the dot product of these contrasts, normalized by the noise covariance:

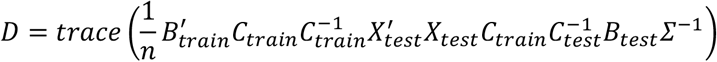

where X_test_ is the design matrix indicating the unique condition of each trial in the test set, C_train_ is the contrast vector the model is trained on, C_test_ the test contrast vector and Σ^−1^ the inverted noise covariance matrix. B_train_ and B_test_ contained the regression parameters of a multivariate general linear model

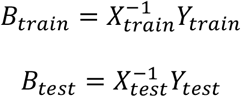

where Y_train_ and Y_test_ are the training and test data sets. The inverted noise covariance matrix Σ^−1^ was estimated using data from a baseline timepoint (−0.5s with respect to the onset of the first cue):

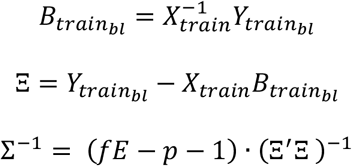

with fE being the degrees of freedom and p the number of sources used. Ξ was regularized towards the unity matrix using a regularization parameter of 0.05.

Because the design matrix and contrast vector include all unique conditions, i.e. all combinations of variable levels (Fig. S1), cvMANOVA independently quantifies information about each variable of interest, while not being confounded by information about the other, potentially correlated variables. In other words, cvMANOVA quantifies the pattern distinctness explained by each variable after discounting the patterns explained by all other variables included in the model.

While cvMANOVA technically constitutes an encoding framework – modelling data variability due to experimental variables – it shares many similarities with commonly used multivariate decoding methods (Hebart and Baker, 2018). Notably, cvMANOVA uses out-of-sample cross-validation to provide a measure of the information contained in neural data about the variables of interest. These estimates can, in principle, also be used to decode experimental variables on individual trials. Due to this close relationship, and to highlight the link to the extensive multivariate decoding literature, we often refer to our results as decoding results.

#### Cross-decoding

To achieve cross-condition decoding, we constructed contrast vectors C_train_ and C_test_ to only contain the conditions to be trained or tested on, respectively. We applied this to estimate neural information within and across the two task conditions (pre and post), as well as the two confidence levels, and the two choices. Additionally, we also used a model trained on all trials, and tested it separately on correct and incorrect trials. To estimate whether information was shared between timepoints, we computed the pattern distinctness when using regression parameters B_train_ from one time point, and B_test_ from another. We repeated this for every pair of time points. In order to assess whether two variables shared a common representational space, we used cross-variable decoding. We implemented this by using a training contrast C_train_ differentiating between the levels of one variable, and a test contrast C_test_ differentiating between the levels of another. Before further processing, all decoding timecourses were smoothed using a Hanning window (500 ms, full width at half maximum). Time-time generalization matrices were smoothed using a 2-dimensional, 100 ms Hanning window.

#### Geometric visualization of representational similarity

We reconstructed low-dimensional geometric representations of neural activity in multiple conditions using the decoding results. Decoding- and cross-decoding values between multiple variables define the distances and angles of condition difference vectors. We used these to plot subsets of conditions in 2D-spaces defined by the axes spanned by two variables of interest. For example, in Figure 4G the length of the choice- and response vectors is given by the magnitude of choice- and response information, respectively; the angle between both is given by the cross-decoding between the two variables. The mapping vector reflects the projection of mapping information onto the 2D-space spanned by choice and response information, indicating that mapping is not represented as an interaction between choice and response.

#### Searchlight analysis

We repeated our main analysis in a searchlight fashion, in order to estimate the spatiotemporal distribution of neural information throughout the trial. For each of the 457 sources, we used cvMANOVA on that source as well as its immediate neighbors, including all 3 dipole directions. In order to maintain comparability between sources, we normalized the resulting pattern distinctness values by the square root of the size of the searchlight (Allefeld and Haynes, 2014; Christophel et al., 2018). After averaging over both hemispheres, we split the searchlight decoding results of all 457 sources into four distinct groups (occipital, temporal, central, frontal) and averaged within each of these areas, to show the spatiotemporal dynamics of neural information. To quantify a shift in choice information from sensory to motor areas, we correlated, for each participant, the cortical distribution of choice information during each timepoint with the distribution of stimulus information during stimulus presentation (1.25 s to 3.25 s), and with the distribution of response information during response execution (from 5.5 s). Statistical significance was assessed using one-tailed cluster-permutation tests.

#### Expected cross-decoding

The maximal amount of shared information between two contexts depends on the amount of information available in each individual context. Thus, in order to assess whether two representations are different, the strength of both representations has to be taken into account and compared with the strength of the shared representation. We thus estimated the expected cross-decoding

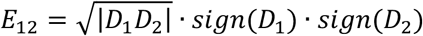

where D_1_ and D_2_ denote the pattern distinctness in the two contexts. The cross-decoding D_12_ between both contexts would be expected to approach E_12_ for identical representations. Any cross-decoding values smaller than E_12_ indicate that the representations are not fully overlapping.

#### Single-trial stimulus prediction

To test whether neural choice representations were informed by the stimulus, we projected each trial’s neural data onto the multivariate axis spanned by yes- and no choices as defined by the cvMANOVA model. We then computed the sign of these single-trial estimates to assess whether it corresponded to the stimulus class.

#### Eye movement control

While we ensured continuous fixation using an online eye movement control at the beginning of each trial, small eye movements can still plausibly confound MEG signals (Quax et al., 2019). We thus repeated our main decoding analysis (Fig. 2) using eye-tracking data. For this purpose, we selected the x-position, y-position, and pupil size signals and averaged them over both eyes. Additionally, we computed the eye position eccentricity as sqrt(x^2^+y^2^). We then applied the same decoding analysis using cvMANOVA, using these 4 channels. We split the 26 participants into the 13 with the highest and lowest choice information in their eye signals, respectively. This revealed that in a subset of participants eye signals were predictive of choice. To test whether this could plausibly explain the neural choice information, we compared the choice decoding timecourses in both splits. As neural choice decoding was, if anything, weaker in those participants with higher choice decoding from the eye signals, the neural decoding was unlikely to be explained by eye movements (Fig. S3).

#### Statistical analysis

We assessed the statistical significance of information using cluster-based sign permutation tests. After determining temporally contiguous clusters during which pattern distinctness was higher than 0 (one-tailed t-test over participants, P < 0.05), we randomly multiplied the information time-course of each participant 10,000 times with either 1 or −1. In each random permutation, we re-computed information clusters and determined the cluster-mass of the strongest cluster. Each original cluster was assigned a p-value by comparing its size to the distribution of sizes of the random permutation’s strongest clusters. The same procedure was used for cross-decoding analyses, however using two-tailed t-tests as true cross-decoding can also be negative. We also tested differences in information using this strategy, namely between response information during “yes” and “no”-choices (Fig. 4E). To test for differences between high- and low-confidence correct and error trials, we averaged data over appropriate time-periods (1.25 to 5.5 s for choice information) and used one-tailed t-tests, as we had a clear unidirectional hypothesis derived from signal detection theory. To determine whether the multivariate patterns underlying two representations were significantly different, we tested whether the empirical cross-decoding was smaller than the expected cross-decoding, again using cluster-based sign permutation tests. Cross-temporal generalization and dynamics were assessed analogously, however using two-dimensional clusters.

#### Software

All analyses were performed in MATLAB, using custom code as well as the Fieldtrip (Oostenveld et al., 2011) and SPM toolboxes. For meta-d’ analyses, we used code from http://www.columbia.edu/~bsm2105/type2sdt/ (Maniscalco and Lau, 2012).

#### Data and materials availability

Preprocessed MEG data and analysis code to reproduce all reported results are available upon reasonable request.

## Acknowledgements

We thank Katrina Quinn for helpful discussions of the manuscript. This study was supported by the European Research Council (ERC) StG 335880 and CoG 864491 (M.S), Deutsche Forschungsgemeinschaft (DFG; German Research Foundation) project 276693517 (SFB 1233) (M.S.) and the Centre for Integrative Neuroscience (DFG, EXC 307) (M.S.). The authors acknowledge support by the state of Baden-Württemberg through bwHPC and the German Research Foundation (DFG) through grant no INST 39/963-1 FUGG (bwForCluster NEMO).

## Supplementary Figures

**Fig. S1.**
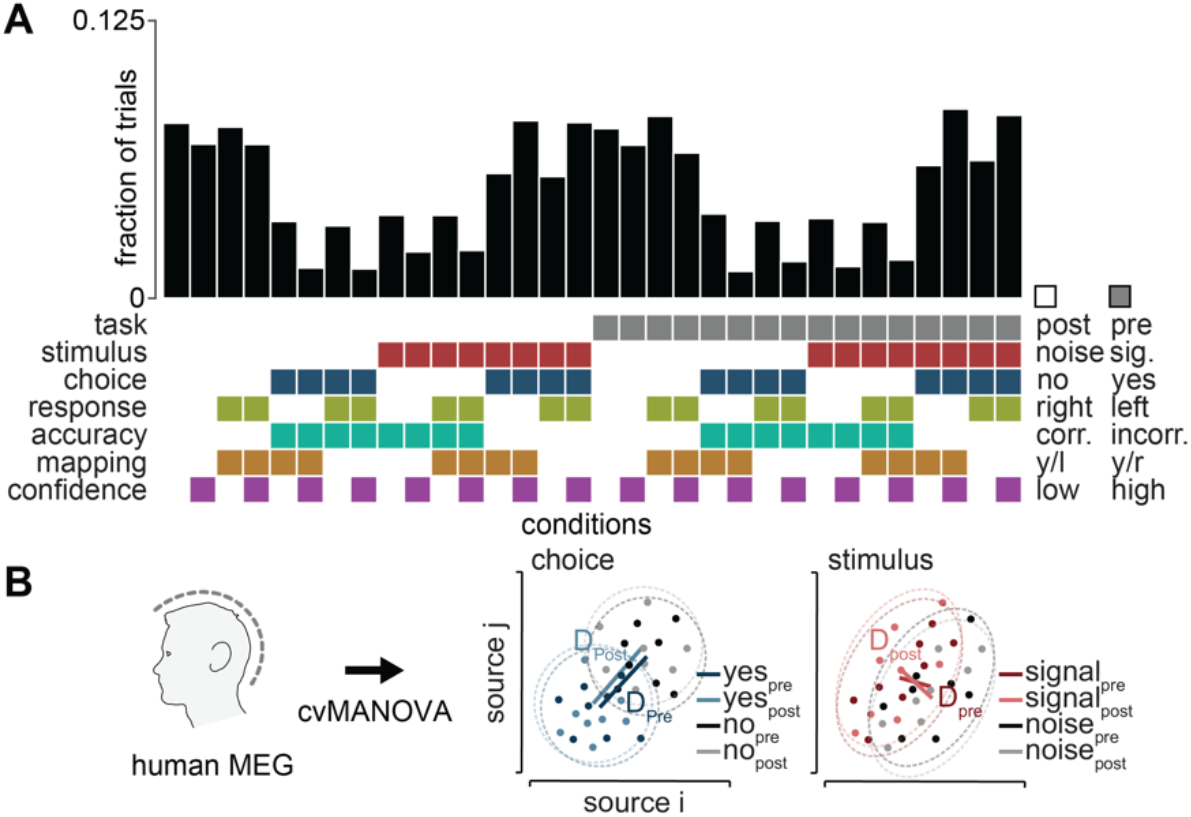
Task variables and analysis schematic. **(A)** Task conditions and behavioral responses. Each combination of the binary task variables constituted one of 32 separate conditions. Because behavioral variables were partially correlated among each other and with experimentally controlled variables (e.g. confidence vs. accuracy and choice vs. stimulus), the number of trials varied across the 32 unique conditions. The rows with variables below the histogram correspond to the contrast vectors employed in the cross-validated MANOVA. **(B)** We performed multivariate pattern analysis using cross-validated MANOVA to estimate the difference D between MEG source level patterns associated with the two levels of each task variable. Importantly, cross-validated MANOVA allowed to independently quantify information about each variable without confounding of other, potentially correlated variables. Each dot represents MEG activity during one of the 32 conditions. For example, choice information is computed as the contrast between all conditions containing “yes”-trials and those containing “no”-trials (middle), stimulus information as the contrast between conditions containing “signal” and “noise” trials (right). For our main analyses, this procedure was applied to the action-linked (“pre”) and action-independent (“post”) contexts separately. Using a cross-decoding framework, the angle between Dpre and Dpost enabled us to assess the degree of similarity between representations of any given variable during action-linked and action-independent contexts.

**Fig. S2.**
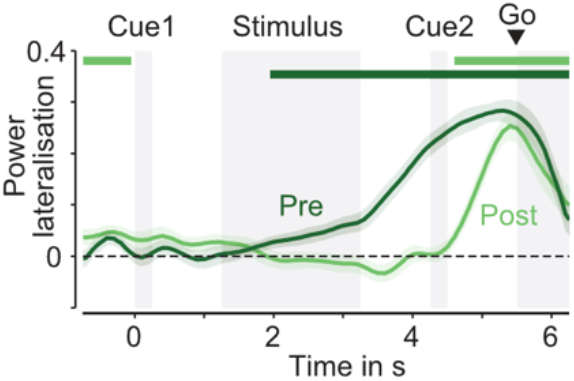
Motor cortical beta lateralization. Beta lateralization predictive of each trial’s response hand was computed for pre and post conditions in an individually localized source in motor cortex for each participant. Horizontal bars indicate significant clusters (cluster permutation, two-tailed, p<0.05). Coloured lines and shaded regions indicate the mean +/- SEM across participants.

**Fig. S3.**
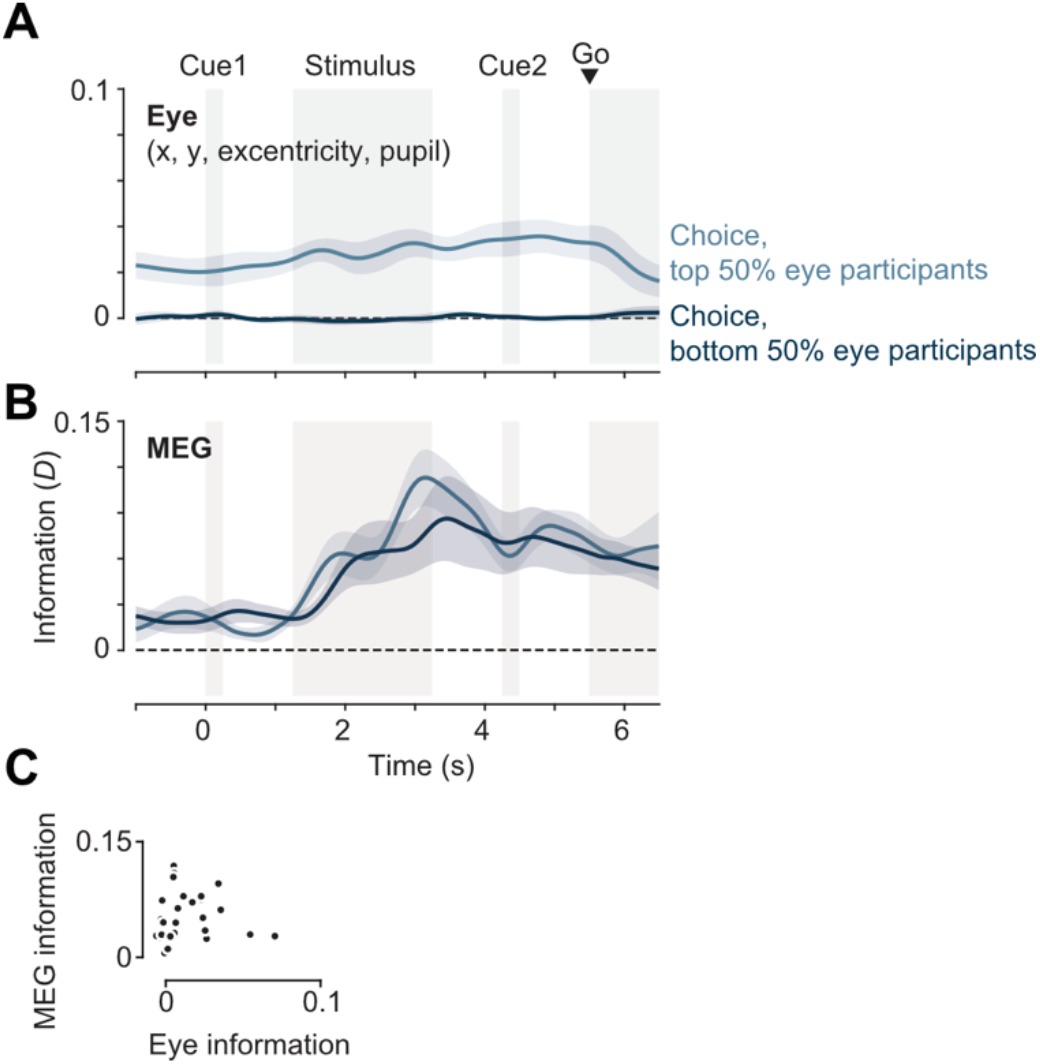
Choice information in eye-tracking data. **(A)** Choice information contained in eye traces (x and y position, eccentricity, and pupil size), split into the 13 participants with the highest decoding values, and the 13 participants with the lowest. Eye traces were thus predictive of choice in a subset of participants. **(B)** Choice information contained in MEG data, split into the same groups as in (A). Participants in whom the eye traces were predictive of choice did not show stronger decoding of choice from MEG data (one-tailed t-test, P > 0.05 for all timepoints). This indicates that choice information in MEG was not driven by eye movements or pupil size. **(C)** There was no positive across-subject relationship between the amount of choice information contained in eye traces and the amount of choice information contained in MEG data. Coloured lines and shaded regions indicate the mean +/- SEM of information across participants.

**Fig. S4.**
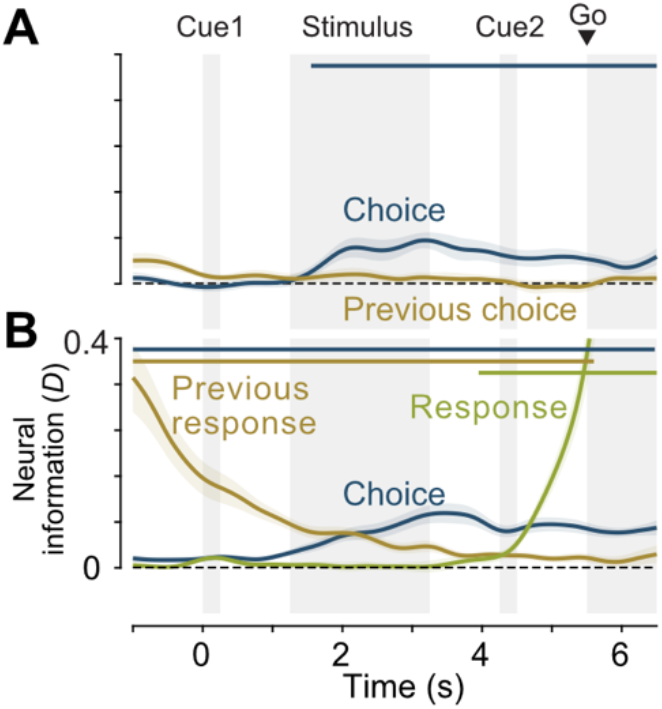
Previous trial information. (A) There was no significant information about the previous choice, and information about the current choice remained significant when including previous choice as a variable. Horizontal lines indicate clusters of significant information (cluster permutation, P < 0.01, N = 23). (B) There was significant information about the previous motor response throughout the trial, but information about the current choice and response remained significant when including previous response as a variable. Horizontal lines indicate clusters of significant information (cluster permutation, P < 0.01, N = 25). Coloured lines and shaded regions indicate the mean +/- SEM of information across participants.

**Fig. S5.**
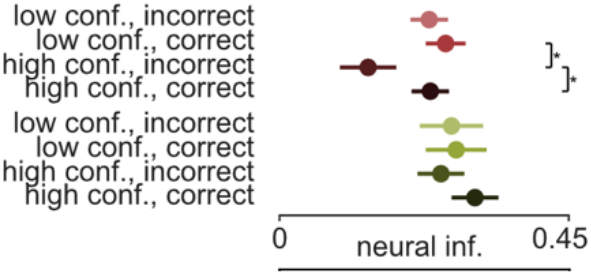
Stimulus- and response representations do not exhibit properties of a decision variable. Time-averaged stimulus information (1.25 to 3.5s) and response information (3.25 to 6.5s) in correct and error, and high- and low confidence trials. The model was trained on both correct and error trials, but trials were split by accuracy for testing. Stars denote significant differences (P < 0.05, two-tailed t-tests, N=19).

